# Inverse Agonist Activity of Angiotensin II Receptor Blocker Is Crucial for Prevention of Aortic Aneurysm Formation in Marfan Syndrome

**DOI:** 10.1101/2022.12.21.521508

**Authors:** Hiroki Yagi, Hiroshi Akazawa, Qing Liu, Masahiko Umei, Hiroshi Kadowaki, Ryo Matsuoka, Akito Shindo, Shun Okamura, Akiko Saga-Kamo, Norifumi Takeda, Issei Komuro

**Affiliations:** Department of Cardiovascular Medicine, Graduate School of Medicine, The University of Tokyo, Tokyo, Japan; Marfan Syndrome Center, The University of Tokyo Hospital, Tokyo, Japan

**Keywords:** ARB, G protein-coupled receptor, inverse agonist, Marfan syndrome, thoracic aortic aneurysm

## Abstract

**Background:** Marfan syndrome (MFS), an inherited disorder caused by *FBN1* gene mutations, causes fatal aortic aneurysm. Selective angiotensin II (Ang II) type 1 (AT_1_) receptor blockade is a preventive option for patients with MFS aortopathy, and recent clinical studies demonstrated that the inhibitory effect of an AT_1_ receptor blocker (ARB) losartan on aortic aneurysm growth is equivalent to that of β-blockers. At present, several ARBs are clinically available, and they have drug-specific differences in pharmacological properties. Especially, inverse agonism of ARBs has potential benefits for cardiovascular protection, but its impact on MFS aortopathy remains poorly understood.

**Methods:** Candesartan-7H is a candesartan derivative which lacks the carboxyl group critical for inverse agonist activity and works as a neutral antagonist for AT_1_ receptor. Candesartan cilexetil (1 mg/kg/day), candesartan-7H (1 mg/kg/day or 20 mg/kg/day), or vehicle was administered to *Fbn1*^C1041G/+^ mice, and aortic aneurysm formation was analyzed using echocardiography, histological staining, and *in situ* MMP assay. Activation of TGF-β signaling and mechanosensitive signaling was studied using western blot and immunohistochemical analysis.

**Results:** Candesartan cilexetil (1 mg/kg/day) and candesartan-7H (20 mg/kg/day) lowered blood pressures equally in *Fbn1*^C1041G/+^ mice, but that candesartan-7H (1 mg/kg/day) did not. Aortic aneurysmal progression in association with aortic wall thickening, degeneration of elastic fibers, deposition of collagen, MMP activation, and TGF-β signaling activation in *Fbn1*^C1041G/+^ mice was significantly suppressed by treatment with candesartan cilexetil (1 mg/kg/day), but not by candesartan-7H even at 20 mg/kg/day. In addition, candesartan cilexetil, but not candesartan-7H, inhibited up-regulation of mechanical stress–responsive transcriptional factor Egr-1 in ascending aorta of *Fbn1*^C1041G/+^ mice.

**Conclusions:** Our findings support a crucial role of inverse agonist activity of ARB for prevention of mechanical stress-induced AT_1_ receptor activation and aortic aneurysm formation in MFS mice.

## Introduction

Marfan syndrome (MFS) is an autosomal dominant connective tissue disorder that affects cardiovascular, skeletal, ocular, and pulmonary systems ^1^. MFS patients are at risk for sudden death in young adulthood due to the development of thoracic aortic aneurysm (TAA) which can progress to aortic dissection or rupture without prior symptoms ^2,3^. Currently, prophylactic aortic root surgery can prolong life expectancy, but there is no sovereign remedy that sufficiently restrains the progression of TAA in MFS patients ^4^.

MFS is caused by mutations in *FBN1* gene, which encodes fibrillin-1 ^5–7^. Fibrillin-1 is a major component of microfibrils within the extracellular matrix (ECM), and also controls the bioavailability of transforming growth factor-β (TGF-β), a potent cytokine that regulates proliferation, differentiation, ECM modeling, and apoptosis ^8^. Several studies using mouse models demonstrated that excessive activation of TGF-β and its downstream signaling pathways, as the consequence of altered structure of fibrillin-1, was a likely culprit underlying TAA progression in MFS ^9,10^. In addition, treatment with angiotensin II (Ang II) type 1 (AT_1_) receptor blockers (ARBs) prevented TAA progression in MFS mice by inhibiting TGF-ß signaling ^11–13^. According to several clinical studies, ARBs (losartan and irbesartan) reduced the rate of aortic root dilatation in MFS patients compared with placebo ^14–16^, but losartan showed no significant benefit over the β-blocker atenolol for the prevention of aortic root dilatation ^17–20^.

The AT_1_ receptor is a seven transmembrane spanning G protein-coupled receptor (GPCR), and is activated upon binding to Ang II ^21,22^. GPCRs are structurally unstable and show significant constitutive activity in an agonist-independent manner ^23,24^. The constitutive activity of wild-type AT_1_ receptor can be detected even in the absence of endogenous expression of Ang II ^25,26^. We and others reported that the AT_1_ receptor can be activated by mechanical stress, independently of Ang II, through a conformational switch of the receptor ^25,27–31^. The agonist-independent constitutive activity or activation of AT_1_ receptor are relevant to the pathogenesis of cardiovascular remodeling ^25,32^, and can be inhibited by inverse agonist which stabilizes inactive conformation of the receptor, but not by antagonist which blocks agonist-dependent receptor activation through competing with agonist binding to the receptor ^23,33^. Therefore, the inverse agonist activity of ARBs provides clinical advantage to inhibit both Ang II-dependent and -independent receptor activation ^34^. The activity of inverse agonism is primarily determined by chemical structure of the drug, and thus is a drug-specific property ^26,29,35–38^.

It was reported that mechanoresponses of vascular smooth muscle cells in MFS mice were impaired ^39^. We hypothesized that mechanical stress-induced activation of AT_1_ receptor contributes to aortic aneurysm progression in *Fbn1*^C1041G/+^ mice. The aim of this study is to determine the impact of inverse agonist activity on the efficacy of ARB to prevent aortic aneurysm formation in MFS mice.

## Methods

### Animal studies

All animal experiments were approved by the Ethics Committee for Animal Experiments of the University of Tokyo (P-14-083 and P-20-040), and adhered strictly to the National Institutes of Health Guide for the Care and Use of Laboratory Animals. Mice were housed in groups of 3–4 animals per cage with a 12 h light-dark cycle at a constant room temperature of 22±1°C with a humidity of 50±15%. The mice were fed a standard chow (CE-2; Clea Japan Inc.) and water *ad libitum* during the experimental period. *Fbn1* ^C1041G/+^ mice were obtained from the Jackson Laboratory ^40,41^, and maintained on a C57BL/6J background. Candesartan cilexetil and candesartan-7H were synthesized by Takeda Pharmaceutical Co., Ltd. Eight-week-old male *Fbn1*^C1041G/+^ mice were treated with candesartan cilexetil (1 mg/kg/day), candesartan-7H (low dose: 1 mg/kg/day, high dose: 20 mg/kg/day), or vehicle in drinking water for 8 weeks ^32^. Mice were sacrificed to collect tissue samples under anesthesia by intraperitoneal injection of sodium pentobarbital (50 mg/kg).

### Transthoracic echocardiography and blood pressure (BP) measurement

Transthoracic echocardiography was performed on mice anesthetized with 2-3% isoflurane inhalation, using a Vevo 2100 system with a 30-MHz probe (FUJIFILM Visualsonics, Inc.) ^42^. For evaluation of aortic diameters, a two-dimensional short-axis view was obtained at the level of aortic root, sinotubular junction (STJ), and ascending aorta at 8 (baseline), 12, and 16 weeks of age. BPs and pulse rates were measured in conscious mice at 16 weeks of age by a tail-cuff method (MK-2000ST NP-NIBP MONITOR; Muromachi Kikai Co., Ltd.) ^42^.

### Western blot analysis

Aortic tissues were harvested, and perivascular adipose tissue was thoroughly removed. The ascending thoracic aortas were crushed by cryo-press, and dissolved in RIPA lysis buffer (50m M Tris-HCl, pH 8.0, 150 mM NaCl, 1% NP-40, 1% SDS, 0.5% Na-deoxycholate, 10 mM Okadaic acid) containing protease inhibitor cocktail (cOmplete ULTRA Tablets Mini EASYpack; Roche Diagnostics). The protein concentration was measured using BCA Protein Assay Kit (Thermo Fisher Scientific). The lysates were mixed with 5x Laemmli Buffer (50 mM Na_2_HPO_4_, 50 mM NaH_2_PO_4_, 2% SDS, 5% 2-mercaptoethanol, 50% glycerol, 0.1% bromophenol blue), boiled at 95°C for 5 min, subjected to SDS–PAGE, and transferred to polyvinylidene difluoride membrane (Merck Millipore). The membranes were blocked with 1% bovine serum albumin (Sigma-Aldrich) or 5% skim milk powder (FUJIFILM Wako Pure Chemical Corporation) in TBST (50 mM Tris-HCl, pH 7.6, 150 mM NaCl, 0.05% Tween 20), and incubated with the following primary antibodies: rabbit monoclonal anti-phosphorylated Smad2 (Ser465/467) antibody (Cell Signaling Technology, #3108), rabbit monoclonal anti-Smad2 antibody (Cell Signaling Technology, #5339), rabbit polyclonal anti-phosphorylated ERK1/2 (Thr202/Tyr204) antibody (Cell Signaling Technology, #9101), rabbit polyclonal anti-ERK1/2 antibody (Cell Signaling Technology, #9102), rabbit monoclonal anti-EGR1 antibody (Cell Signaling Technology, #4154), and rabbit monoclonal anti-GAPDH antibody (Cell Signaling Technology, #2118). The membranes were then incubated with horseradish peroxidase-conjugated anti-mouse (Jackson Immuno Research Laboratories, Inc., #115-035-146) or anti-rabbit IgG antibody (Jackson Immuno Research Laboratories, Inc., # 111-035-003). Immunoreactive signals were detected with ECL Prime Western Blotting Detection Reagent (GE Healthcare Biosciences), and acquired using Lumino Graph I (ATTO Corp.). The integrated density per image area of interest was measured using an NIH Image J software (NIH, Research Branch; http://imagej.nih.gov/ij/).

### Histological and immunohistochemical analysis

The ascending aortas were excised, fixed immediately in 10% neutralized formalin (Sakura Finetek Japan Co., Ltd.), and embedded in paraffin. Serial 5 μm sections were stained with hematoxylin-eosin (HE) for morphological analysis, elastica van Gieson (EVG) for detection of elastic fibers, Masson’s trichrome for detection of collagen fibers, and Alcian blue for staining proteoglycan. Degeneration of elastic fibers was evaluated by calculating the number of elastic fiber breaks per high power field at 4-5 different representative locations of EVG-stained sections, and the numbers were averaged by 2 observers blinded to the genotype and treatment for each mouse ^43^.

For immunohistochemical analysis, deparaffinized sections were rehydrated, boiled to retrieve antigens (10 mM citrate buffer, pH 6, for phosphorylated Smad2, ERK1/2), and blocked with 10% goat or rabbit serum in PBS for 60 min and with Avidin/Biotin Blocking Kit (Vector Laboratories, Inc.) for 15 min. The sections were incubated with the following primary antibodies: rabbit monoclonal anti-phosphorylated Smad2 antibody (Ser465/467) (Cell Signaling Technology, #3108), rabbit polyclonal anti-phosphorylated ERK1/2 antibody (Thr202/Tyr 204) (Cell Signaling Technology, #9101). The Vectastain ABC kit (Vector Laboratories, Inc.) and DAB Peroxidase Substrate Kit (Vector Laboratories, Inc.) were used to detect the primary antibodies. The sections were counterstained with hematoxylin and mounted in Mount-Quick (Daido Sangyo Co., Ltd.).

### *In situ* matrix metalloproteinase (MMP) activity assay

*In situ* MMP assay was performed on freshly frozen OCT-embedded sections (5 μm) of ascending aortas. Fluorescein-conjugated DQ Gelatin from Pig Skin (50 μg/ml, Thermo Fisher Scientific) in a reaction buffer (0.5 M Tris-HCl, pH 7.6, 1.5 M NaCl, 50 mM CaCl2, 2 mM NaN3) was applied to the sections, and incubated at 37°C for 24 h. Sections for negative control were incubated in the presence of 5 mM EDTA. The sections were mounted in ProLong Gold Antifade Reagent with DAPI (Thermo Fisher Scientific). An all-in-one fluorescence microscope (BZ-9000; Keyence) was used to detect gelatinase (MMP-2 and MMP-9) activity as green fluorescence.

### Statistical analysis

All data are expressed as means ± SEM. For two-group comparisons, a two-tailed Student’s *t*-test was used. For multiple comparisons, one-way ANOVA with Tukey’s multiple comparisons test was used for group comparisons. Values of *P* < 0.05 were considered statistically significant. All data were statistically analyzed using GraphPad PRISM software version 8.0 (GraphPad Software).

## Results

### Aneurysm formation in *Fbn1*^C1041G/+^ mice is suppressed by treatment with an inverse agonist, but not antagonist, for AT_1_ receptor

We previously reported that candesartan suppressed mechanical stretch–induced helical movement of AT_1_ receptor as an inverse agonist, and thereby inhibited mechanical stress-induced AT_1_ receptor activation and pressure overload-induced cardiac hypertrophy even in *angiotensinogen-null* mice ^25,27^. The tight binding between the carboxyl group at the benzimidazole ring of candesartan and specific residues of the AT_1_ receptor is critical for the potent inverse agonist activity of candesartan to inhibit Ang II-independent receptor activation ^27,34,36^. A candesartan derivative (candesartan-7H), lacking the carboxyl group critical for inverse agonist activity, inhibits Ang II-dependent AT_1_ receptor activation as a neutral antagonist, but cannot inhibit Ang II-independent constitutive activity of AT_1_ receptor or mechanical stress-induced AT_1_ receptor activation ^27,32,34,36^.

We previously reported that treatment with candesartan-7H (high dose: 20 mg/kg/day) suppressed Ang II-induced BP elevation in C57BL/6 male mice, almost equally as treatment with candesartan cilexetil (1 mg/kg/day) did ^32^. However, treatment with candesartan-7H (low dose: 1 mg/kg/day) had almost no inhibitory effect on Ang II-induced BP elevation ^32^.

We first treated *Fbn1*^C1041G/+^ mice at 8 weeks of age with vehicle, candesartan cilexetil (1 mg/kg/day), candesartan-7H (1 mg/kg/day), or candesartan-7H (20 mg/kg/day) for 8 weeks. Tail-cuff measurements revealed that candesartan cilexetil (1 mg/kg/day) and candesartan-7H (20 mg/kg/day) equally lowered both systolic and diastolic BP in *Fbn1*^C1041G/+^ mice, but that candesartan-7H (1 mg/kg/day) did not show BP-lowering effect (**Figure 1A**). Pulse rate and body weight were comparable among the groups (**Figure 1 A and 1B**).

**Figure 1.**
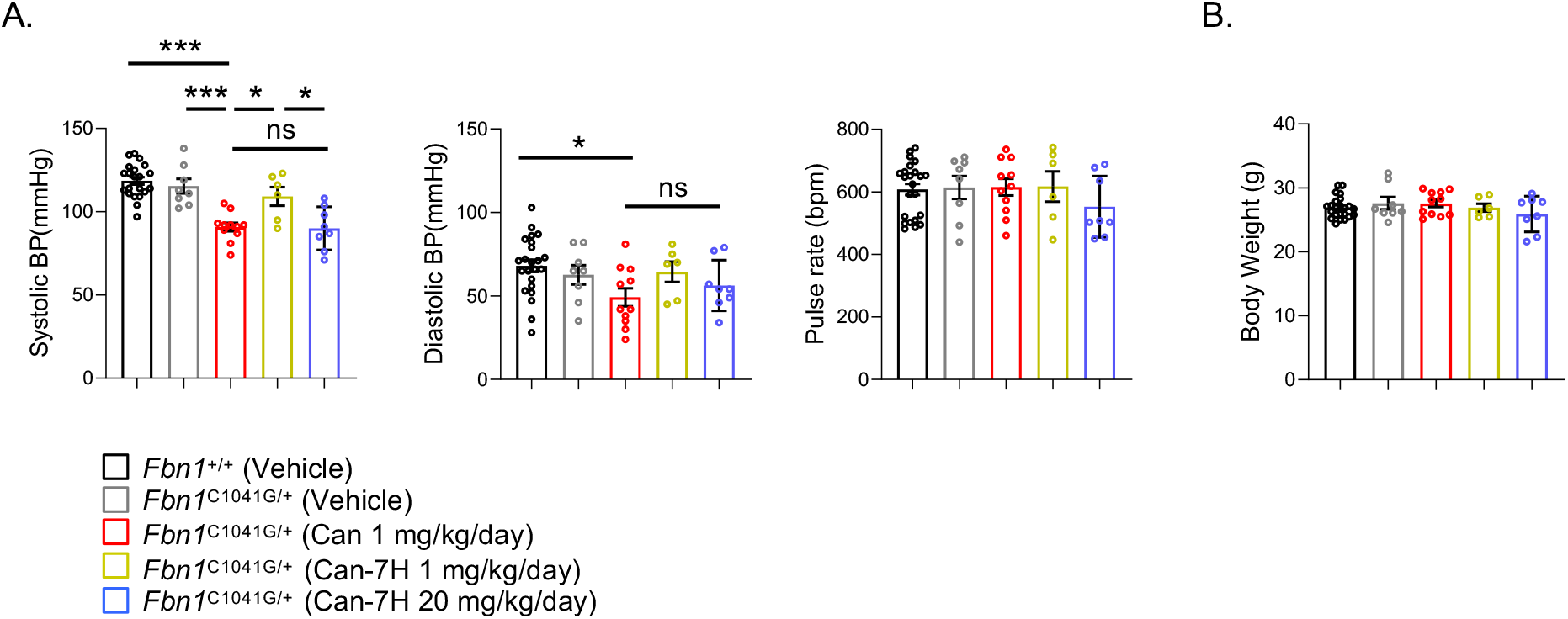
Blood pressure (BP) lowering effects of candesartan cilexetil and candesartan-7H in *Fbn1*^C1041G/+^ mice. (A) The systolic and diastolic BP, and pulse rate of 16-week-old vehicle-treated *Fbn1^+/+^*mice (n = 22) and *Fbn1*^C1041G/+^ mice treated with vehicle (n = 8), candesartan cilexetil (Can) (1 mg/kg/day) (n = 11), candesartan-7H (Can-7H) (1 mg/kg/day) (n = 6), or Can-7H (20 mg/kg/day) (n = 8). The data are mean ± SEM. \**P* < 0.05, \*\*\**P* < 0.001, ns, not significant, one-way ANOVA with Tukey’s multiple comparisons test. (B) Body weight of 16-week-old vehicle-treated *Fbn1^+/+^* mice (n = 23) and *Fbn1*^C1041G/+^mice treated with vehicle (n = 8), Can (1 mg/kg/day) (n = 11), Can-7H (1 mg/kg/day) (n = 6), or Can-7H (20 mg/kg/day) (n = 8). The data are mean ± SEM. One-way ANOVA with Tukey’s multiple comparisons test.

We next measured the diameters of aortic root, STJ, and ascending aorta at baseline and 8 weeks after treatment. Notably, candesartan cilexetil almost completely prevented progression of aortic dilatation at all levels of aortic root, STJ, and ascending aorta in *Fbn1*^C1041G/+^ mice (**Figure 2**). However, candesartan-7H, even at a high dose (20 mg/kg/day), did not significantly prevent progression of aortic dilatation (**Figure 2**). Histologically, candesartan cilexetil significantly attenuated aortic wall thickening, degeneration of elastic fibers, deposition of collagen, and MMP activation in ascending aorta of *Fbn1*^C1041G/+^ mice, as compared with treatment with vehicle or candesartan-7H (low dose and high dose) (**Figure 3**). These results suggest that inverse agonist activity of candesartan is crucial for inhibition of aneurysm formation in *Fbn1*^C1041G/+^ mice.

**Figure 2.**
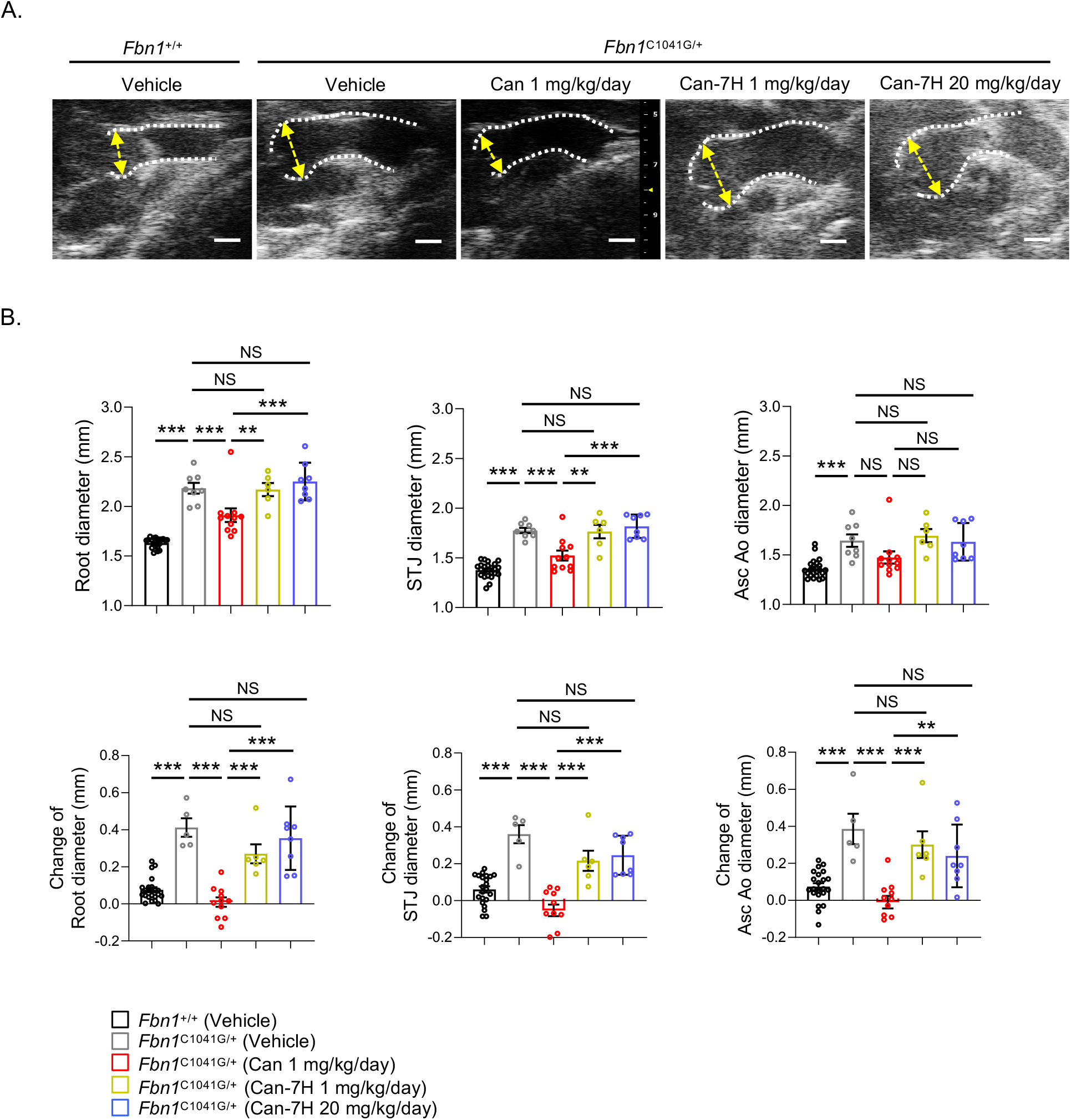
Prevention of aneurysm formation in *Fbn1*^C1041G/+^mice by candesartan cilexetil, but not by candesartan-7H. (A) Representative ultrasound images of ascending aorta in 16-week-old vehicle-treated *Fbn1^+/+^* mice and *Fbn1*^C1041G/+^ mice treated with vehicle, candesartan cilexetil (Can) (1 mg/kg/day), candesartan-7H (Can-7H) (1 mg/kg/day), or Can-7H (20 mg/kg/day). Arrowheads indicate aortic root. Scale bar, 1 mm. (B) Ascending aorta diameters at three different levels (Root, aortic root; STJ, sinotubular junction; Asc Ao, ascending aorta) in 16-week-old vehicle-treated *Fbn1^+/+^* mice (n = 23) and *Fbn1*^C1041G/+^ mice treated with vehicle (n = 8), Can (1 mg/kg/day) (n = 11), Can-7H (1 mg/kg/day) (n = 6), or Can-7H (20 mg/kg/day) (n = 8) (upper panels). Changes of ascending aorta diameters at Root, STJ, and Asc Ao in vehicle-treated *Fbn1^+/+^* mice (n = 22) and *Fbn1*^C1041G/+^ mice treated with vehicle (n = 5), Can (1 mg/kg/day) (n = 11), Can-7H (1 mg/kg/day) (n = 6), or Can-7H (20 mg/kg/day) (n = 8) after 8 weeks of treatment (lower panels). The data are mean ± SEM. \*\**P* < 0.01, \*\*\**P* < 0.001, ns, not significant, one-way ANOVA with Tukey’s multiple comparisons test.

**Figure 3.**
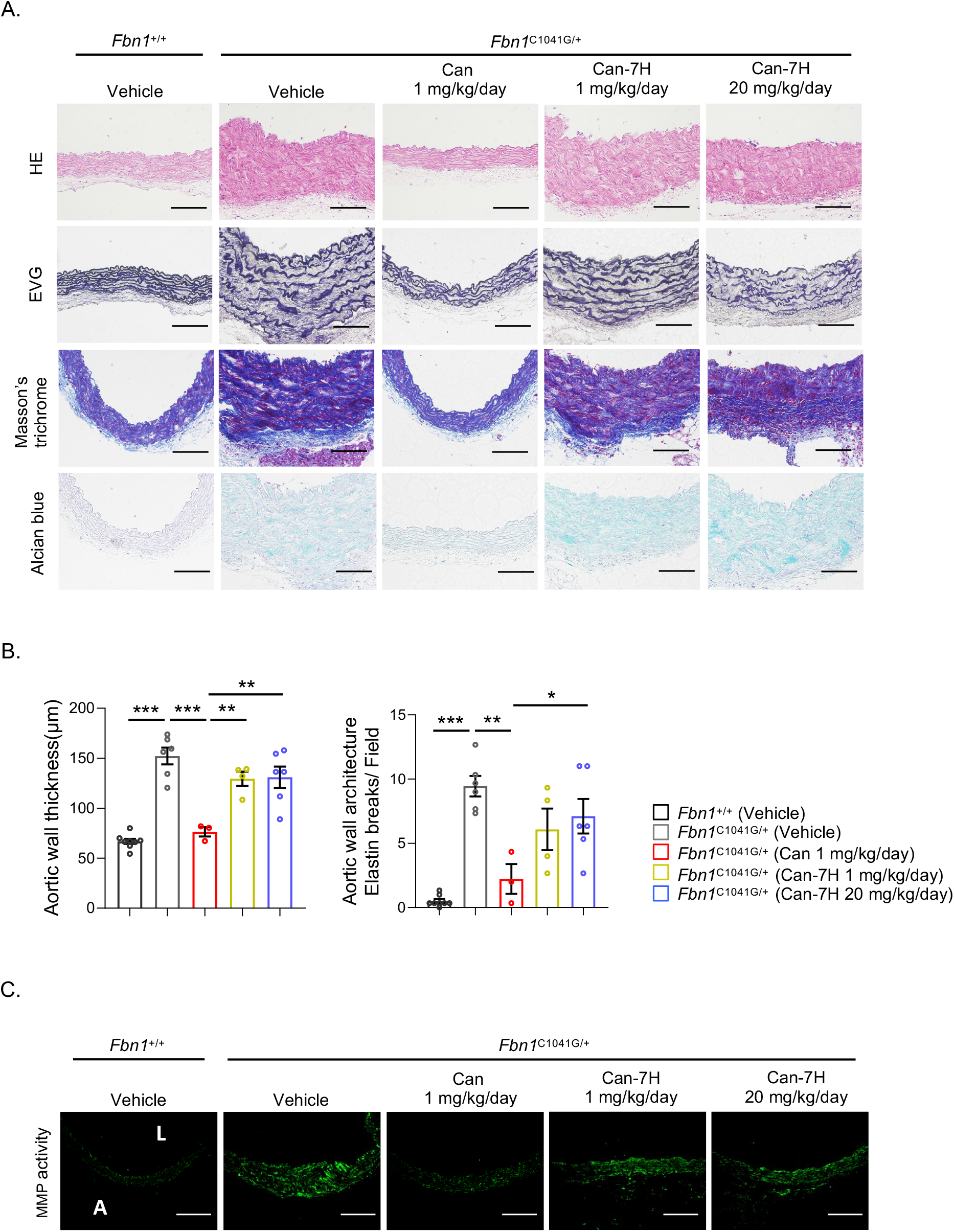
Prevention of histopathological changes in ascending aorta of *Fbn1*^C1041G/+^ mice by candesartan cilexetil, but not by candesartan-7H. (A) Histological analysis with hematoxylin-eosin (HE) staining, elastica van Geison (EVG) staining, Masson’s trichrome staining, and Alcian blue staining in ascending aorta of 16-week-old vehicle-treated *Fbn1*^+/+^ mice and *Fbn1*^C1041G/+^ mice treated with vehicle, candesartan cilexetil (Can) (1 mg/kg/day), candesartan-7H (Can-7H) (1 mg/kg/day), or Can-7H (20 mg/kg/day). Scale bar, 100 μm. (B) Measurement of aortic wall thickness and aortic architecture indicated by the number of breaks of the elastic fiber per field in 16-week-old vehicle-treated *Fbn1^+/+^* mice (n =8) and *Fbn1*^C1041G/+^ mice treated with vehicle (n = 6), Can (1 mg/kg/day) (n = 3), Can-7H (1 mg/kg/day) (n = 4), or Can-7H (20 mg/kg/day) (n = 6). The data are mean ± SEM. \**P* < 0.05, \*\**P* < 0.01, \*\*\**P* < 0.001, one-way ANOVA with Tukey’s multiple comparisons test. (C) *In situ* zymography for gelatinase activity in ascending aorta of 16-week-old vehicle-treated *Fbn1^+/+^* mice and *Fbn1*^C1041G/+^ mice treated with vehicle, Can (1 mg/kg/day), Can-7H (1 mg/kg/day), or Can-7H (20 mg/kg/day). Scale bars, 100 μm.

### Activation of TGF-ß signaling in ascending aorta of *Fbn1*^C1041G/+^ mice is suppressed by treatment with an inverse agonist, but not antagonist, for AT_1_ receptor

Recent studies demonstrated that TGF-ß signaling not only via the canonical (Smad-dependent) pathway but also via the non-canonical (Smad-independent) pathway was involved in aortic aneurysm formation in *Fbn1*^C1041G/+^ mice ^9–12^. Both immunohistochemical analysis and western blot analysis revealed that phosphorylation levels of Smad2 and ERK1/2 in ascending aorta of *Fbn1*^C1041G/+^ mice were significantly attenuated by treatment with candesartan cilexetil, but not by treatment with vehicle or candesartan-7H (low dose and high dose) (**Figure 4**). These results suggest that inverse agonist activity of candesartan is indispensable for inhibition of both canonical and non-canonical TGF-β signaling activation in the ascending aorta of *Fbn1*^C1041G/+^mice.

**Figure 4.**
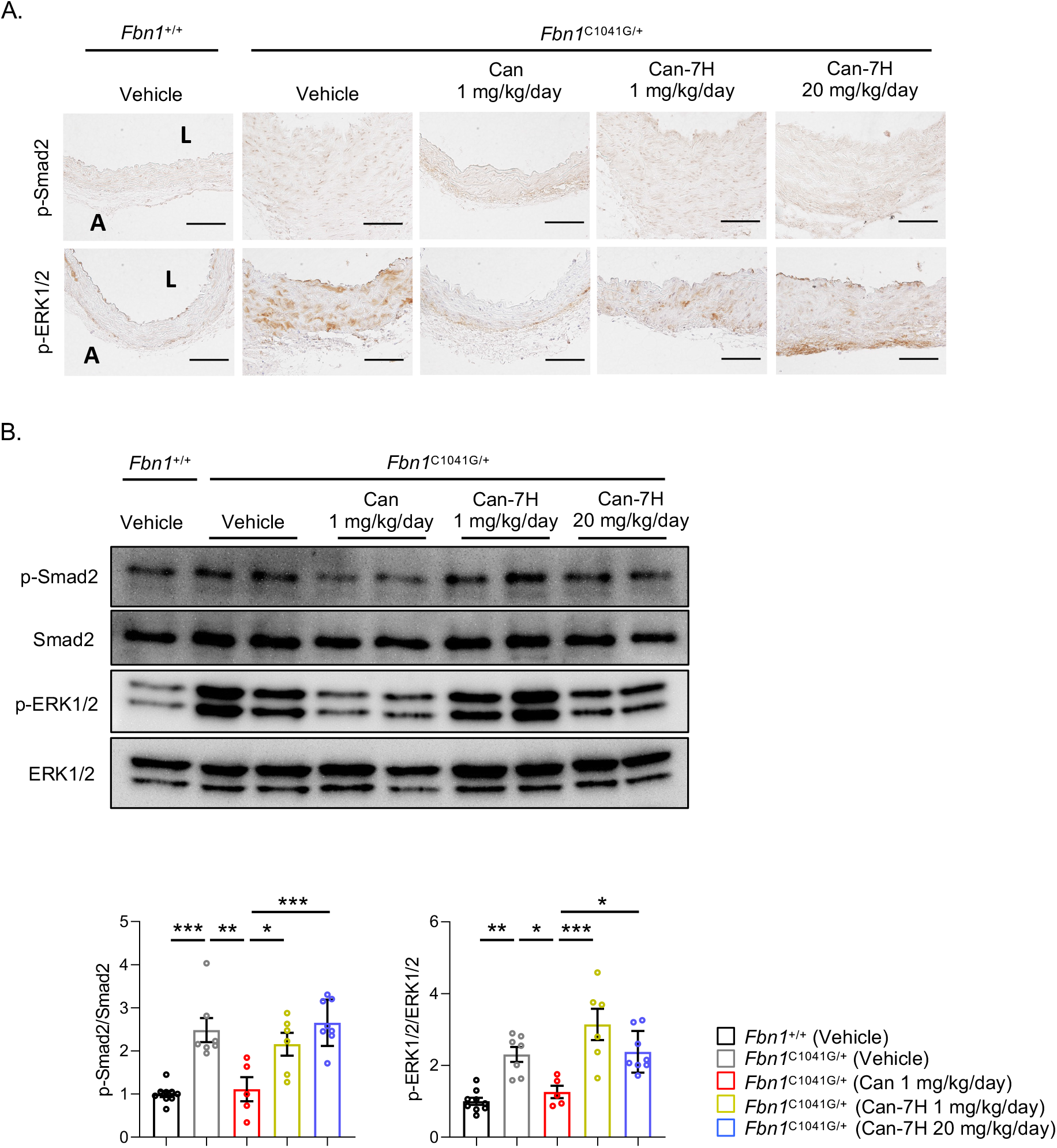
Prevention of TGF-ß signaling activation in ascending aorta of *Fbn1*^C1041G/+^ mice by candesartan cilexetil, but not by candesartan-7H. (A) Immunostaining for phosphorylated Smad2 (p-Smad2) and phosphorylated ERK1/2 (p-ERK1/2) in ascending aorta of 16-week-old vehicle-treated *Fbn1^+/+^* mice and *Fbn1*^C1041G/+^ mice treated with vehicle, candesartan cilexetil (Can) (1 mg/kg/day), candesartan-7H (Can-7H) (1 mg/kg/day), or Can-7H (20 mg/kg/day). Scale bars, 100 μm. (B) Immunoblot analysis of p-Smad2, Smad2, p-ERK1/2, ERK1/2 in ascending aorta of 16-week-old vehicle-treated *Fbn1^+/+^* mice and *Fbn1*^C1041G/+^ mice treated with vehicle, Can (1 mg/kg/day), Can-7H (1 mg/kg/day), or Can-7H (20 mg/kg/day). The intensity of each band was quantified by densitometric analysis, and quantitation graphs for the p-Smad2/Smad2 (n = 5–9) and p-ERK1/2/ERK1/2 (n = 5–9) are shown (mean ± SEM). \**P* < 0.05, \*\**P* < 0.01, \*\*\**P* < 0.001, one-way ANOVA with Tukey’s multiple comparisons test.

### Up-regulation of mechanosensitive Egr-1 protein in ascending aorta of *Fbn1*^C1041G/+^mice is suppressed by treatment with an inverse agonist, but not antagonist, for AT_1_ receptor

We previously demonstrated that mechanosensitive signaling was aberrantly activated in ascending aorta of *Fbn1*^C1041G/+^ mice and MFS patients ^42^. The expression levels of early growth response-1 (Egr-1) protein, a mechanical stress–responsive transcriptional factor ^44–46^ were significantly increased in ascending aorta of *Fbn1*^C1041G/+^ mice, as compared with *Fbn1^+/+^* mice (**Figure 5**). Interestingly, treatment with candesartan cilexetil suppressed the up-regulation of Egr-1 protein in ascending aorta of *Fbn1*^C1041G/+^mice, but treatment with vehicle or candesartan-7H (low dose and high dose) did not (**Figure 5**). These results suggest that inverse agonist activity of candesartan is essential for inhibition of aberrant activation of mechanical stress-induced signaling in the ascending aorta of *Fbn1*^C1041G/+^ mice.

**Figure 5.**
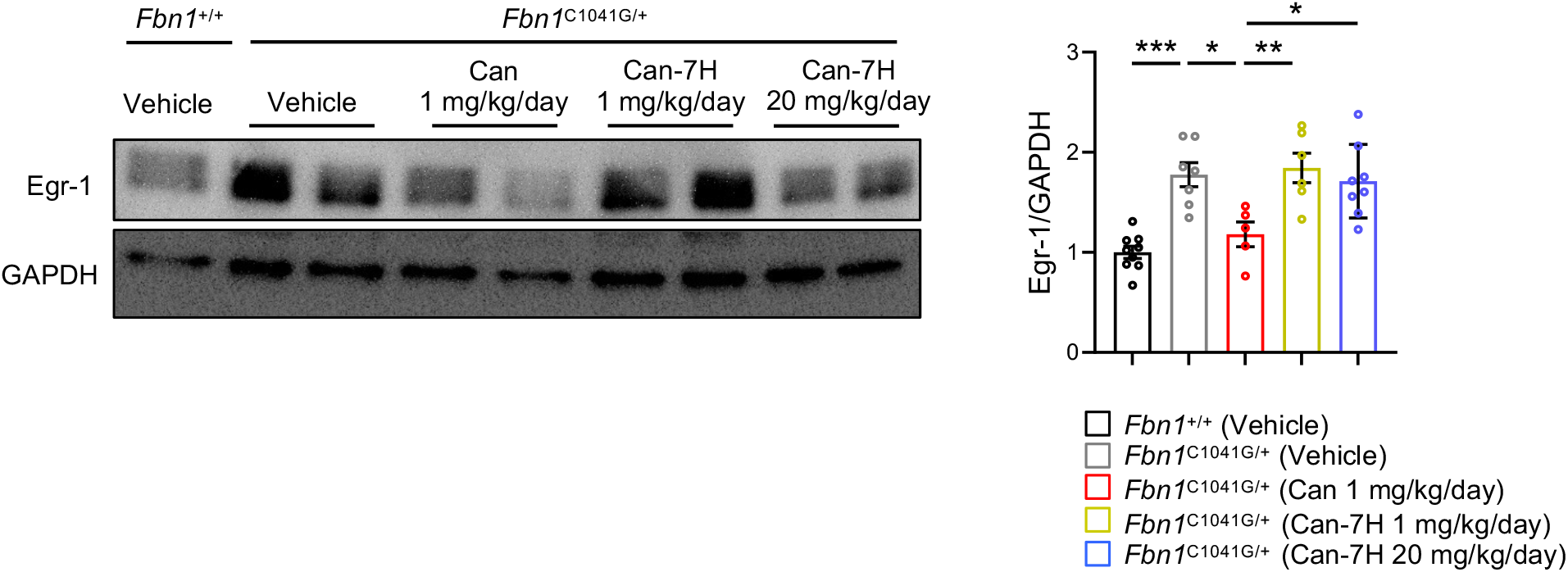
Prevention of mechano-sensitive Egr-1 protein up-regulation in ascending aorta of *Fbn1*^C1041G/+^ mice by candesartan cilexetil, but not by candesartan-7H. Immunoblot analysis of Egr-1 protein in ascending aorta of 16-week-old vehicle-treated *Fbn1^+/+^* mice and *Fbn1*^C1041G/+^ mice treated with vehicle, candesartan cilexetil (Can) (1 mg/kg/day), candesartan-7H (Can-7H) (1 mg/kg/day), or Can-7H (20 mg/kg/day). The intensity of each band was quantified by densitometric analysis and corrected for the amount of Gapdh protein as an internal control (n = 5–9) (mean ± SEM). \**P* < 0.05, *\*\*P* < 0.01, *\*\*\*P* < 0.001, one-way ANOVA with Tukey’s multiple comparisons test.

## Discussion

In this study, we demonstrated that candesartan cilexetil (1 mg/kg/day), but not candesartan-7H even at a higher dose (20 mg/kg/day), prevented aortic root dilatation, in association with histopathological changes, MMP activation, TGF-ß signaling activation, and up-regulation of mechanosensitive Egr-1 protein in ascending aorta of *Fbn1*^C1041G/+^ mice. Candesartan cilexetil (1 mg/kg/day) and candesartan-7H (20 mg/kg/day) equally lowered BP in *Fbn1*^C1041G/+^ mice, suggesting that candesartan-7H can inhibit Ang II-dependent AT_1_ receptor activation as a neutral antagonist at a higher dose (20 mg/kg/day), as previously described ^32^. Candesartan-7H lacks the carboxyl group of candesartan critical for inverse agonist activity, and cannot inhibit mechanical stress-induced AT_1_ receptor activation ^27,32,34,36^. Therefore, our study highlights inverse agonist activity as a crucial pharmacological property of ARBs to prevent the progression of TAA in *Fbn1*^C1041G/+^ mice.

The role of the AT_1_ receptor in the pathogenesis of MFS aortopathy has been extensively investigated using murine MFS models. Indeed, losartan suppressed aortic root dilatation in *Fbn1*^C1041G/+^ mice ^11^, but the preventive effect of losartan was not fully observed when Ang II type 2 (AT_2_) receptor was knocked out ^13^. Although administration of antisense oligonucleotide against angiotensinogen attenuated aortic root dilatation in *Fbn1*^C1041G/+^ mice ^47^, an angiotensin-converting enzyme inhibitor enalapril, which inhibits Ang II formation and blocks both AT_1_ and AT_2_ receptor signaling, did not sufficiently inhibit aortic root dilatation, despite showing BP-lowering effects similar to those of losartan ^13^. This indicates that the beneficial effects of losartan require activation of AT_2_ receptor signaling. However, direct stimulation of AT_2_ receptor with an AT_2_ receptor agonist Compound 21 was ineffective for prevention of aortic root dilatation in *Fbn1*^C1041G/+^ mice ^48^. A previous study also attributed the beneficial effects of losartan to an AT_1_ receptor-independent protection of endothelial function ^49^. On the other hand, sequential measurement of aortic dimensions revealed that genetic knockout of AT_1_ receptor attenuated aortic root dilatation in *Fbn1*^C1041G/+^ mice ^47^, indicating that AT_1_ receptor activation are crucial for the development of MFS aortopathy.

However, it is still an open question how AT_1_ receptor is activated in ascending aorta of MFS, because there has been no evidence of local activation of the renin-angiotensin system. A previous study using aortic tissue of MFS patients showed a significant decrease in AT_1_ receptor expression and a significant increase in AT_2_ receptor expression, with a significant higher Ang II concentration ^50^. Importantly, transcriptomic analysis revealed that the components of renin-angiotensin system were not changed in ascending aorta of *Fbn1*^C1041G/+^ mice ^47^. AT_1_ receptor is expressed in multiple cell types within aortic tissue, including endothelial cells, vascular smooth muscle cells, fibroblasts, and inflammatory cells ^21,22^. Cell type-specific knockout of AT_1_ receptor revealed the dominant role of endothelial AT_1_ receptor in the pathogenesis of MFS aortopathy in mice. Endothelial cell-specific AT_1_ receptor knockout modestly suppressed aortic dilatation and increased survival, whereas smooth muscle cell-specific AT_1_ receptor knockout did not show apparent changes in TAA pathology in fibrillin-1 hypomorphic mice (*Fbn1*^mgR/mgR^) ^51^. Although it is not fully determined which cell type is most responsible for AT_1_ receptor-mediated pathogenesis and how AT_1_ receptor is activated, one possible mechanism underlying increased AT_1_ receptor signaling in MFS aorta is mechanical stress-induced activation of AT_1_ receptor in aortic endothelium.

Mechanical stress is an important trigger to promote aortic dilatation in MFS, as evidenced by both *in vitro* and *in vivo* experiments ^46^. Traction force microscopy revealed that force-generating capacity of aortic vascular smooth muscle cells of *Fbn1*^mg△lpn/+^mice was impaired with disorganization of actin stress fiber network and focal adhesions, indicating mechanoresponse impairment in aortic wall of MFS mice ^39^. It was also reported that passive mechanical myocardial tension was reduced in *Fbn1*^mgR/mgR^ mice ^52^. Importantly, the cardiomyopathy phenotype in *Fbn1*^mgR/mgR^ mice was restored by genetic knockout of AT_1_ receptor or treatment with losartan, but not by genetic knockout of angiotensinogen nor treatment with enalapril ^52^. In vascular smooth muscle cells differentiated from MFS patient-derived induced pluripotent stem cells, cyclic stretch increased activation of mechanosensitive ß_1_-integrin and p38 mitogen-activated protein kinase, leading to exacerbation of MFS phenotype and promotion of apoptosis ^53^. It is likely that altered structure and function of fibrillin-1 and inadequate extracellular matrix remodeling may change the stiffness of aortic tissue and perturb normal responses to mechanical forces ^46^, leading to cellular alterations characteristic of MFS aortopathy. Consistently, acute pressure overload by transverse aortic constriction in wild-type mice was sufficient to induce an aortic smooth muscle cell phenotype, similar to that of *Fbn1*^C1041G/+^ mice ^54^.

Therefore, it is likely that activation of AT_1_ receptor signaling in MFS aorta can be most efficiently prevented by ARBs with potent inverse agonist activity, not by AT_1_ receptor antagonists nor by angiotensin-converting enzyme inhibitors, because inverse agonist for AT_1_ receptor can inhibit Ang II-independent and mechanical stress-induced receptor activation as well as Ang II-dependent receptor activation ^34,36^. Given this background, we demonstrated, in this study, that the loss of inverse agonist activity abolished therapeutic efficacy of candesartan to prevent aortic aneurysm formation in *Fbn1*^C1041G/+^ mice. The potency of inverse agonist activity differs among ARBs according to each chemical structure ^36^. Most of ARBs share a common biphenyltetrazole ring structure, and the unique side chain structures contribute to specific drug-receptor bindings that determine inverse agonist activity to stabilize the receptor in an inactive conformation ^36^. We previously reported that the bindings of the carboxyl group at the benzimidazole ring of candesartan to specific residues of AT_1_ receptor are responsible for the potent inverse agonism in inhibiting mechanical stretch-induced activation of AT_1_ receptor ^27^. Similarly, drug-receptor interactions involving the carboxyl group and hydroxyl group of olmesartan were essential for potent inverse agonism on stretch-induced AT_1_ receptor activation ^29^. Although the inverse agonism of commercially available ARBs has not been fully determined, valsartan, irbesartan, azilsartan, and EXP3174 (active metabolite of losartan) also reduce the constitutive GTPase stimulating activity of AT_1_ mutant receptor, while losartan does not ^26,35,38,55,56^. According to a recent study, a subpressor dose of valsartan attenuated aortic dilatation in *Fbn1*^C1041G/+^ mice, to a similar degree as a pressor dose of losartan. Pharmacological properties including inverse agonist activity may contribute to the difference in therapeutic efficacy between valsartan and losartan ^57^. From a view point of inverse agonist activity, head-to-head comparisons of ARBs in preclinical studies will provide a clue to understand the drug effect of ARBs on the prevention of aortic aneurysm formation in MFS.

In conclusion, inverse agonist activity of ARB is crucial for prevention of mechanical stress-induced AT_1_ receptor activation and aortic aneurysm formation in MFS mice. Therefore, we suggest that ARBs with potent inverse agonist activity will have superior therapeutic efficacy. A proof-of-concept implementation will achieve pathophysiology-based optimal medication to restrain the progression of TAA and improve the prognosis and quality of life in MFS patients.

## Acknowledgments

None

## Nonstandard Abbreviations and Acronyms

Ang II: angiotensin II
ARB: angiotensin II type 1 receptor blocker
Asc Ao: ascending aorta
AT_1_ receptor: angiotensin II type 1 receptor
BP: blood pressure
ECM: extracellular matrix
Egr-1: early growth response-1
ERK: extracellular signal-regulated kinase
EVG: elastica van Gieson
GPCR: G protein coupled receptor
HE: hematoxylin-eosin
MFS: Marfan syndrome
STJ: sinotubular junction
TAA: thoracic aortic aneurysm
TGF-β: transforming growth factor-β

## Highlights

- Candesartan cilexetil (inverse agonist for angiotensin II receptor) (1 mg/kg/day) and candesartan-7H (candesartan derivative lacking the carboxyl group critical for inverse agonism) (20 mg/kg/day) equally lower blood pressure in *Fbn1*^C1041G/+^ mice.
- Aortic aneurysm formation in *Fbn1*^C1041G/+^ mice is suppressed by treatment with candesartan cilexetil, but not by candesartan-7H.
- Up-regulation of mechanosensitive Egr-1 protein in ascending aorta of *Fbn1*^C1041G/+^mice is suppressed by treatment with candesartan cilexetil, but not by candesartan-7H.

